# Association Rule Mining of the Human Gut Microbiome

**DOI:** 10.1101/2022.11.27.518104

**Authors:** Yiyan Zhang, Shanlin Ke, Xu-Wen Wang, Yang-Yu Liu

**Affiliations:** Department of Biostatistics, Harvard T.H. Chan School of Public Health, Boston, Massachusetts 02115, USA; Channing Division of Network Medicine, Department of Medicine, Brigham and Women’s Hospital and Harvard Medical School, Boston, Massachusetts 02115, USA; Center for Artificial Intelligence and Modeling, The Carl R. Woese Institute of Genomic Biology, University of Illinois at Urbana-Champaign, Champaign, IL 61820, USA

## Abstract

The human gut carries a vast and diverse microbial community that is essential for human health. Understanding the structure of this complex community requires quantitative approaches. Traditional co-occurrence and correlation analyses typically focus on pair-wise relationships and ignore higher-order relationships. Association rule mining (ARM) is a well-developed technique in data mining and has been applied to human microbiome data to identify higher-order relationships. Yet, existing attempts suffer from small sample sizes and low taxonomic resolution. Here we leverage the curatedMetagenomic Database (CMD) to resolve those issues. We first infer association rules from gut microbiome samples of a large cohort of healthy individuals (n=2,815) in CMD. Then we compare those rules with that inferred from samples of individuals with different diseases: Inflammatory Bowel Disease (IBD, n=768), Colorectal cancer (CRC, n=368), Impaired Glucose Tolerance (IGT, n=199), and Type 2 Diabetes (T2D, n=164). Finally, we demonstrate that using ARM as a feature selection tool can improve the performance of microbiome-based disease classification. Together, this study provides a comprehensive study of higher-order microbial relationships in the human gut microbiome and highlights the importance of incorporating association rules in microbiome-based disease classification.

## INTRODUCTION

The human body (especially the gastrointestinal tract) harbors a diverse of microorganisms that includes bacteria, protozoa, archaea, viruses, and fungi (Huseyin et al., 2017). The gut microbiota has notable influence on the host health and the dysbiosis of gut microbiota has been associated with many diseases (Thursby & Juge, 2017). Instead of a simple collection of microorganisms, the gut microbiota is a complex community, where different microorganisms interact with each other in different ways (e.g., nutrient competition, metabolic cross-feeding, etc.) and result in complex relationships (Hooper et al., 2012; Koskella et al., 2017).

Microbial co-occurrence networks have been employed to explore the structure of microbial communities. In a microbial co-occurrence network, nodes represent microbes and edges represent their co-occurrence relationship (Ma et al., 2017). Exploring microbial co-occurrence patterns has shown the growing potential in medical applications, e.g., in the study of the pathogenesis and clinical treatment of colorectal diseases (Baxter et al., 2016). Numerous methods have been developed to construct microbial co-occurrence networks. Those methods typically focus on pairwise co-occurrence patterns. Yet, there is mounting evidence that higher-order co-occurrence of microbial species may be as significant as individual species (Gould et al., 2018).

Association rule mining (ARM) is a rule-based machine learning method for discovering interesting patterns between variables in large databases. It was initially introduced to discover regularities or patterns between products (or items) in large-scale transaction data (Agrawal et al., 1993; Piatetsky-Shapiro, 1991). Recently, ARM has been proven as a promising technique to explore higher-order microbial co-occurrence patterns (Naulaerts et al., 2015; Tandon et al., 2016). The basic idea is that we can treat each microbiome sample as a “transaction” and the present species are “items”. Hence, traditional ARM tools can be directly applied to analyze microbiome data. Yet, previous studies typically leveraged 16S rRNA gene sequencing datasets with limited resolution on taxonomy profiling (Giulia et al., 2022; M. Liu et al., 2021; Tandon et al., 2016). Moreover, the small sample sizes used in those studies raised two issues: (1) one has to collapse taxa into higher taxonomic level (e.g., genus or even family level) to mine association rules; (2) the association rules identified from small cohorts might be spurious.

To resolve those issues, we first improved the standard ARM framework to eliminate spurious association rules and preserve interpretability. Then we applied the improved ARM to a large-scale metagenomics database --- the curatedMetagenomicData to explore the higher-order occurrence patterns of the human gut microbiome at the species level. We inferred association rules from gut microbiome samples of a large cohort of healthy individuals (n=2,815) in CMD, representing an unprecedentedly large study on the association rules mining of human gut microbiome. We found that microbial species involved in butyrate producing dominate the co-occurrence patterns in the healthy gut microbiome. Then we compared those baseline association rules inferred from samples of healthy individuals with that inferred from samples of individuals with different diseases: Inflammatory Bowel Disease (IBD, n=768), Colorectal cancer (CRC, n=368), Impaired Glucose Tolerance (IGT, n=199), and Type 2 Diabetes (T2D, n=164). In particular, by comparing the higher-order co-occurrence patterns of the obesity cohort and the healthy cohort, we found the difference between the frequent co-occurrence sets were due to species that have been associated with obesity in literature. Finally, we employed ARM as a feature selection tool in microbiome-based disease classification, finding that ARM-based feature selection can improve the classification performance for all the four diseases and four standard classifiers.

## RESULTS

### Database

In this study, we leveraged gut microbiome samples from a publicly available large-scale metagenomic database --- curatedMetagenomicData (CMD; version 3.6.0) which contains uniformly processed human microbiome taxonomic abundances data and phenotypic data (Pasolli et al., 2017). The species-level bacterial, fungal, and archaeal taxonomic abundances for each sample were profiled with MetaPhlAn3 (Beghini et al., n.d.). We selected samples based on two criteria: the host age is between 18 and 65; no antibiotic use. We selected samples from healthy individuals, as well individuals with one of the following diseases: Inflammatory Bowel Disease (IBD), Colorectal cancer (CRC), Impaired Glucose Tolerance (IGT), and Type 2 Diabetes (T2D). The final dataset includes 2,815 healthy samples, 768 IBD samples, 368 CRC samples, 199 IGT samples, and 164 T2D samples. We filtered out those species whose prevalence were lower than 0.1, resulting in 200 species.

### Association rule mining (ARM) framework

Consider a set of microbiome samples represented by an abundance table 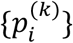, where 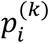 is the relative abundance of species-*i* in sample-*k* (**Fig.1a**). We transform the continuous relative abundances to binary presence/absence variables, i.e., 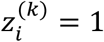 (or 0) if 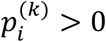 (or = 0) in sample-*k*, to form the “transaction table” (**Fig.1b**). Then, we randomly selected 75% of samples (“transactions”) to form a “subsample transaction table” (**Fig.1c**).

**Figure 1.**
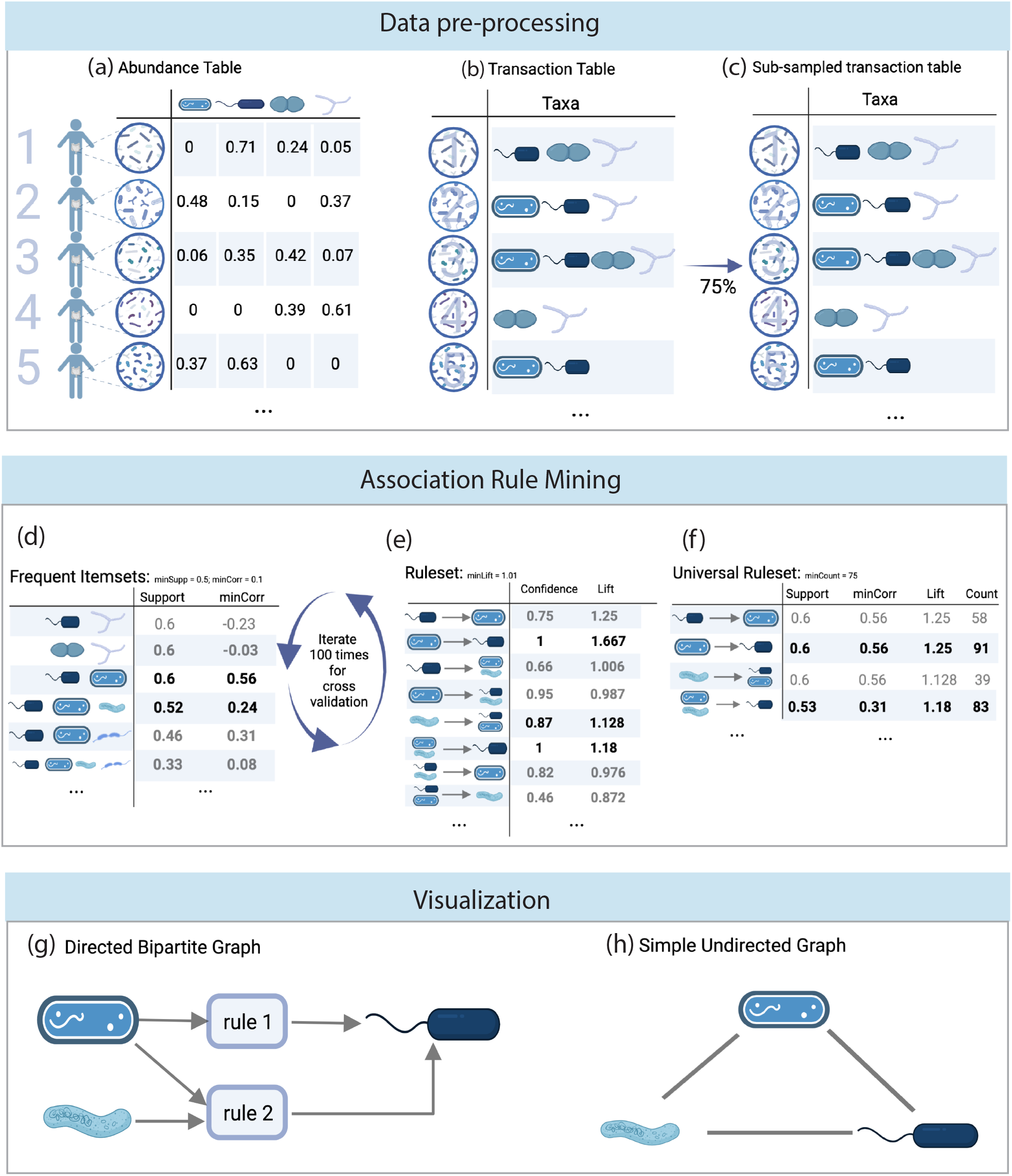
Schematic workflow of association rule mining (ARM) for microbiome data. (**Top**) Data pre-processing: We transform the relative abundance table (**a**) into the presence/absence or transaction table (**b**). Then we randomly sample 75% of samples to form a subsampled transaction table (**c**). (**Middle**) Association rule mining: We apply the Frequent Pattern Growth (FP-growth) algorithm to the subsampled transaction table to find frequent itemsets (**d**). For those frequent itemsets with support values above our support threshold (0.7), we compute pair-wise correlations of within each frequent itemset based on the transaction table, and drop those itemsets with minimum correlation minCorr(*X*) < 0.1. From the remaining frequent itemsets, we infer association rules with lift > 1.01, resulting in the candidate rule set (**e**). The traditional “support-confidence” framework would be prone to the rules which have high-prevalence rhs. In order to accommodate this issue, we consider lift as an the rule measurement threshold which can avoid false high quality rules. We iterate 100 times of the rule mining process and only consider the rules occurred more than 75 times. This results in the universal rule set (**f**). (**Bottom**) Visualization: To visualize those universal rules, we can plot a directed bipartite graph, connecting rules to their species with directed edges connecting LHS species nodes to rule modes and then to RHS species nodes (**g**). For a rule of length *n*, we can also construct a *n*-clique for each length n rule showing the co-occurrence patterns **(h)**.

Traditional ARM composes of two steps: *frequent itemset generation* and *association rule inference* (Agrawal et al., 1993). For frequent itemset generation, we extract all patterns of co-occurrence of items, i.e., itemsets, with the frequency exceeding the *Support* threshold using the Frequent Pattern Growth (FP-growth) algorithm. Here, the *Support* of an itemset *X*, denoted as supp(*X*), is the proportion of transactions (i.e., microbiome samples in our context) in which the itemset *X* (i.e., a set of microbial species) is present. Note that supp(*X*) naturally depends on the prevalence of species in *X*. To set a reasonable *Support* threshold, we examined the prevalence of each species, finding that most prevalent species are from phylum of *Firmicutes, Bacteroidetes*, and *Actinobacteria*, and a considerable number of species (33% for Health cohort, 19% for IBD, 33% for CRC, 28.5% for IGT, and 35% for T2D) are present in more than half of samples. Hence, we set the *Support* threshold as 0.7, and we filter out those frequent itemsets with supp(*X*) < 0.7. To accommodate the unique features of microbiome data, we use the minimum pair-wise correlation to further filter out frequent itemsets. In particular, for each frequent itemset *X* generated by the FP-growth algorithm, we compute the Pearson correlation coefficient of the abundances of each species pair within *X* across all the samples in the subsampled transaction table) and denote the minimum correlation coefficient among all the species pairs in *X* as minCorr (*X*). We filter out those frequent itemsets with minCorr(*X*) < 0.1 (**Fig.1d**). For association rule inference, we identify the association rules (in the forms of *X* ⇒ *Y*, meaning the presence of the LHS itemset *X* implies the presence of the RHS itemset *Y*) from the frequent itemsets with conditional probability exceeding certain threshold. All possible combinations of the items in a frequent itemset are arranged into each side of the rule, and the conditional probability is computed to evaluate the quality of the rule. For a rule *X* ⇒ *Y*, one quality measure is its *Confidence*, defined as conf(*X* ⇒ *Y*) = supp(X ∩ *Y*)/supp(*X*), i.e., the fraction of all transactions containing *X* that also contain *Y*. Another quality measure is *Lift*, defined as lift(*X* ⇒ *Y*) = supp(X ∩ *Y*)/[supp(*X*) × supp(*Y*)]. For microbiome data, the *Confidence* measure tend to infer rules which have high prevalence of *Y*. In order to eliminate those spurious rules, we use *Lift* to infer association rules. In particular, we filter out those rules with lift(*X* ⇒ *Y*) < 1.01, resulting a candidate rule set (**Fig.1e**). To further eliminate spurious rules, we repeat the above process 100 times to randomly generate 100 subsample transaction tables (“realizations”), and filter out those rules present in less than 75 realizations, resulting a universal rule set (**Fig.1f**).

To visualize the association rules, we construct a directed bipartite graph connecting rules to species involved in the rules (**Fig.1g**). Directed edges are added to indicate LHS species *X* and RHS species *Y* of any given rule *X* ⇒ *Y*. In order to conduct further comprehensive network analysis, for a rule of length *n* (i.e., involving *n* species), we can also construct a simple undirected graph, i.e., a *n*-clique (**Fig.1h**).

### Association rules in healthy gut microbiome

We first applied the ARM framework (as depicted in Fig.1) to the 2,815 healthy gut microbiome samples in CMD. We identified 5,230 universal association rules. There are 25 species involved in all the universal rules. The longest rules contain the co-occurrence information of up to 6 species, and we have 62 such rules. The shortest rules still contain the co-occurrence information of 2 species, and we have 16 rules of length 2. The highest confidence among all rules is 0.9996, and the highest *Lift* among all rules is 1.1975.

The top-20 rules sorted based on their *Lift* values were visualized as a directed bipartite graph (see **Fig.2**). The rule with the highest *Lift* is {*F. prausnitzii, D. longicatena, A. hallii*} }⟹ {*F. saccharivorans, D. formicigenerans*}.

**Figure 2.**
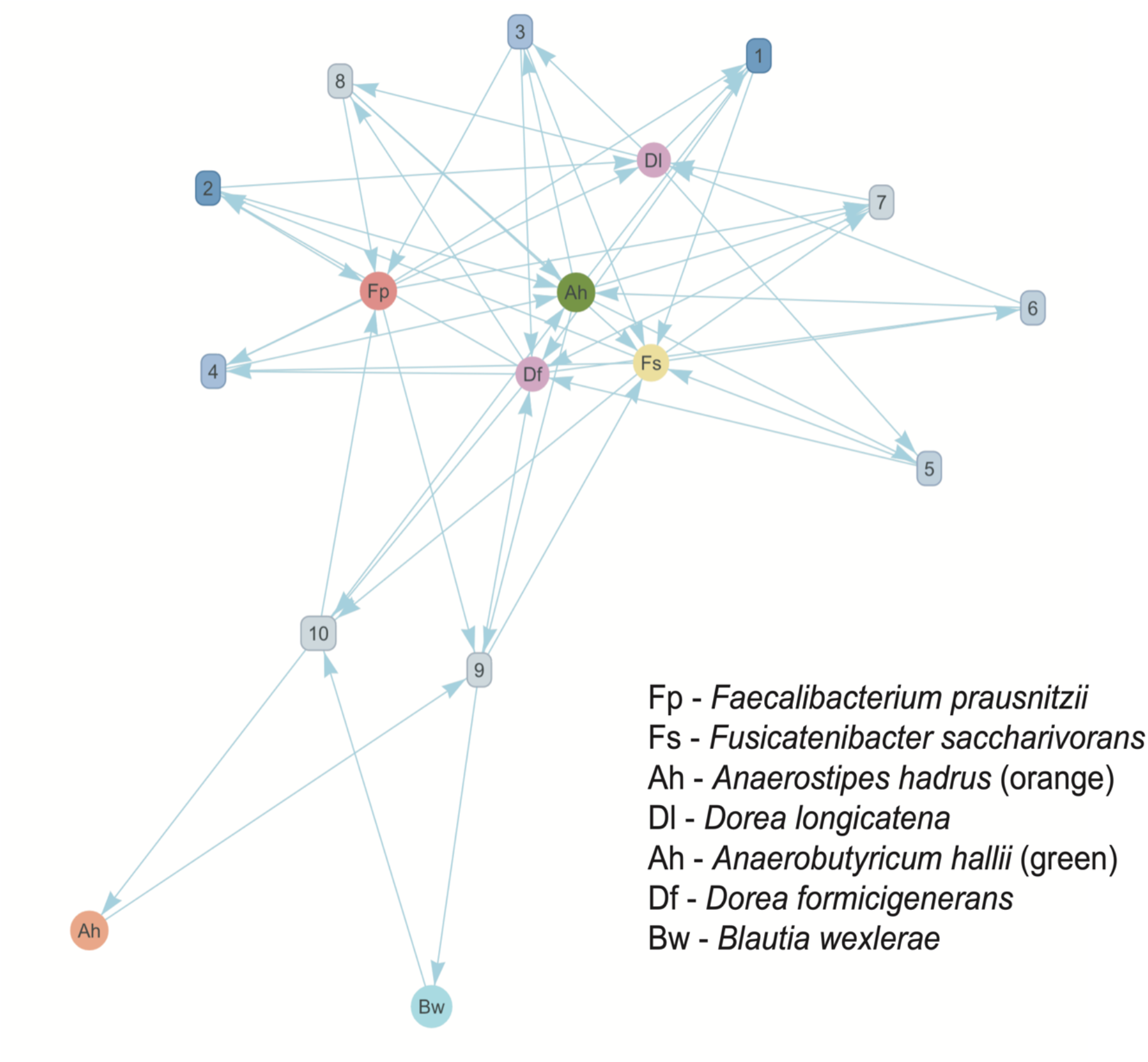
Top-10 association rules inferred from healthy gut microbiome samples. Circle nodes represent species colored by their genus, and species node size is proportional to the prevalence of the species. Square nodes represent association rules shaded by their *Lift* values.

### Association rules in diseased gut microbiome

Following the same procedure, we also inferred the association rules of diseased gut microbiome. In particular, we analyzed 768 IBD samples, 368 CRC samples, 199 IGT samples, and 164 T2D samples. We found 110,118 association rules for IBD, 2,276 rules for CRC; 8,638 rules for IGT; and 81,660 rules for T2D. The maximum rule lengths are 8, 5, 6, and 8 for IBD, CRC, IGT, and T2D, respectively. The highest *Lift* from each rule set is 1.1680 for IBD, 1.1783 for CRC, 1.2010 for IGT, and 1.2037 for T2D. The species involved in the rulesets for IBD, CRC, IGT, and T2D are 31, 28, 32, and 32 respectively.

To compare the association rules mined from healthy and diseased gut microbiome, we first compared their frequent itemsets. By plotting frequent itemsets as incidence matrices, we observed a noticeable nested structure for all the five groups: Healthy, IBD, CRC, IGT, and T2D. In other words, those species belonging to smaller frequent itemsets tend to belong to bigger frequent itemsets (**Fig.3**). We calculated the nestedness value based on the classical nestedness metric based on overlap and decreasing fill (NODF) (Almeida-Neto et al., 2008), finding NODF_Healthy_ = 21.9457%, NODF_IBD_ = 25.9190%, NODF_CRC_ = 15.3155%, NODF_IGT_ = 15.5949%, and NODF_T2D_ = 25.7004%.

**Figure 3.**
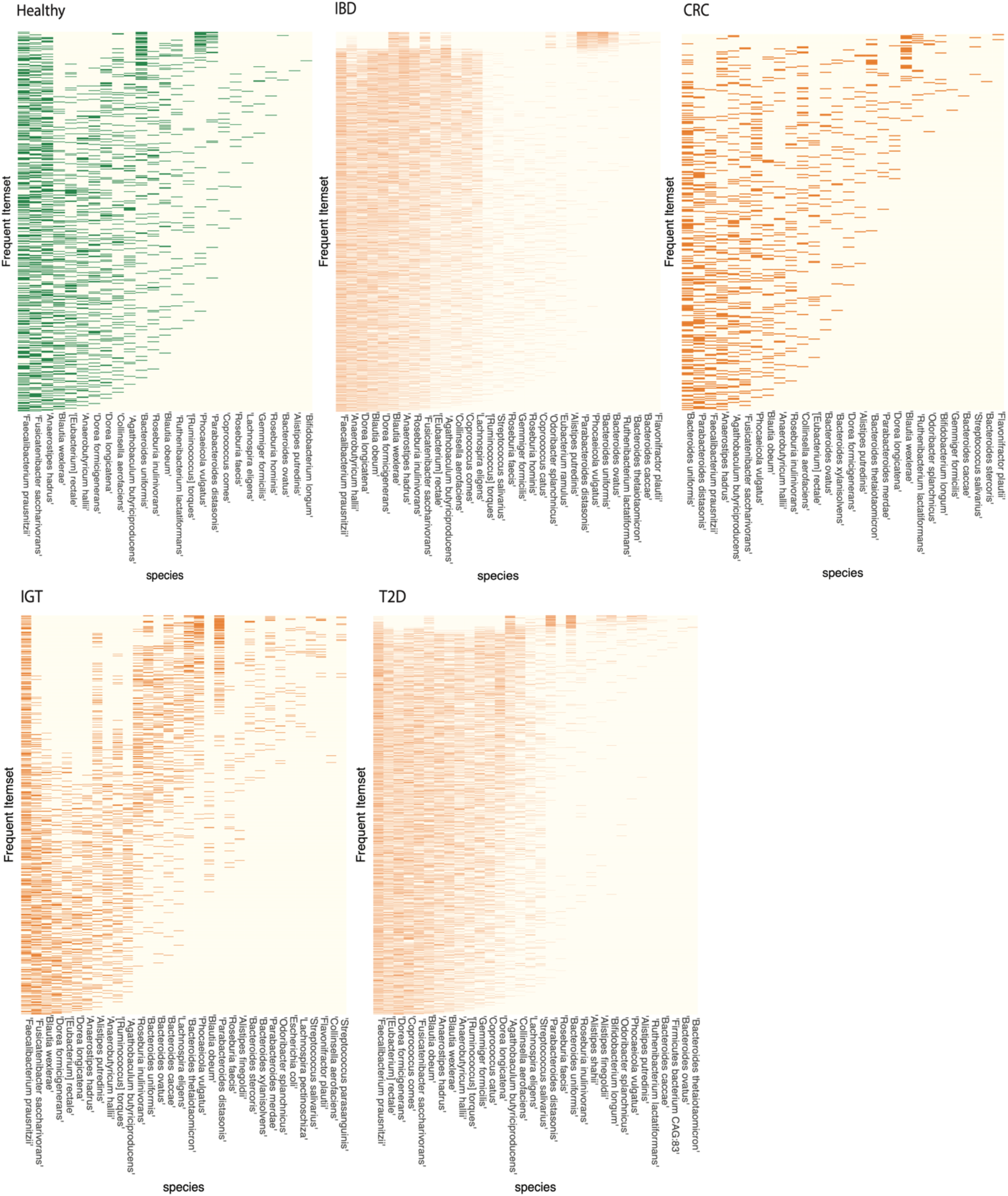
Frequent itemsets generated from healthy and diseased gut microbiome samples. IBD: Inflammatory Bowel Disease, CRC: Colorectal cancer, IGT: Impaired Glucose Tolerance, T2D: Type 2 Diabetes. Each frequent itemset is represented by an incidence matrix, where the presence of a species in a frequent itemset is colored either in green (for Healthy controls) or in orange (for diseased samples). We organized those incidence matrices using the Nestedness Temperature Calculator to emphasize their nested structure. The nestedness value of each matrix is calculated based on the classical NODF measure: NODF_Healthy_ = 21.9457%, NODF_IBD_ = 25.9190%, NODF_CRC_ = 15.3155%, NODF_IGT_ = 15.5949%, and NODF_T2D_ = 25.7004%.

We then directly compared the association rules inferred from of healthy and diseased microbiome. We consider three types of rules with decreasing level of similarity: (1) *common rules* that are shared by both groups: healthy and diseased microbiome; (2) *differentially composed rules*, which are themselves unique to one group, but both the LHS and RHS itemsets are shared by both groups; (3) *unique rules*, which are completely unique to one group --- neither the LHS nor the RHS item sets appear in the rule set of another group. We categorized and summarized the similarity between healthy and each disease cohort in terms of their association rules (see **Table. 3**). Some of the rules in the third case can be used to differentiate between the healthy and disease samples which we will discuss later. For the first two cases with shared LHS and RHS list from both rule sets, we can directly visualize the rule sets (see **Fig.4**).

**Figure 4.**
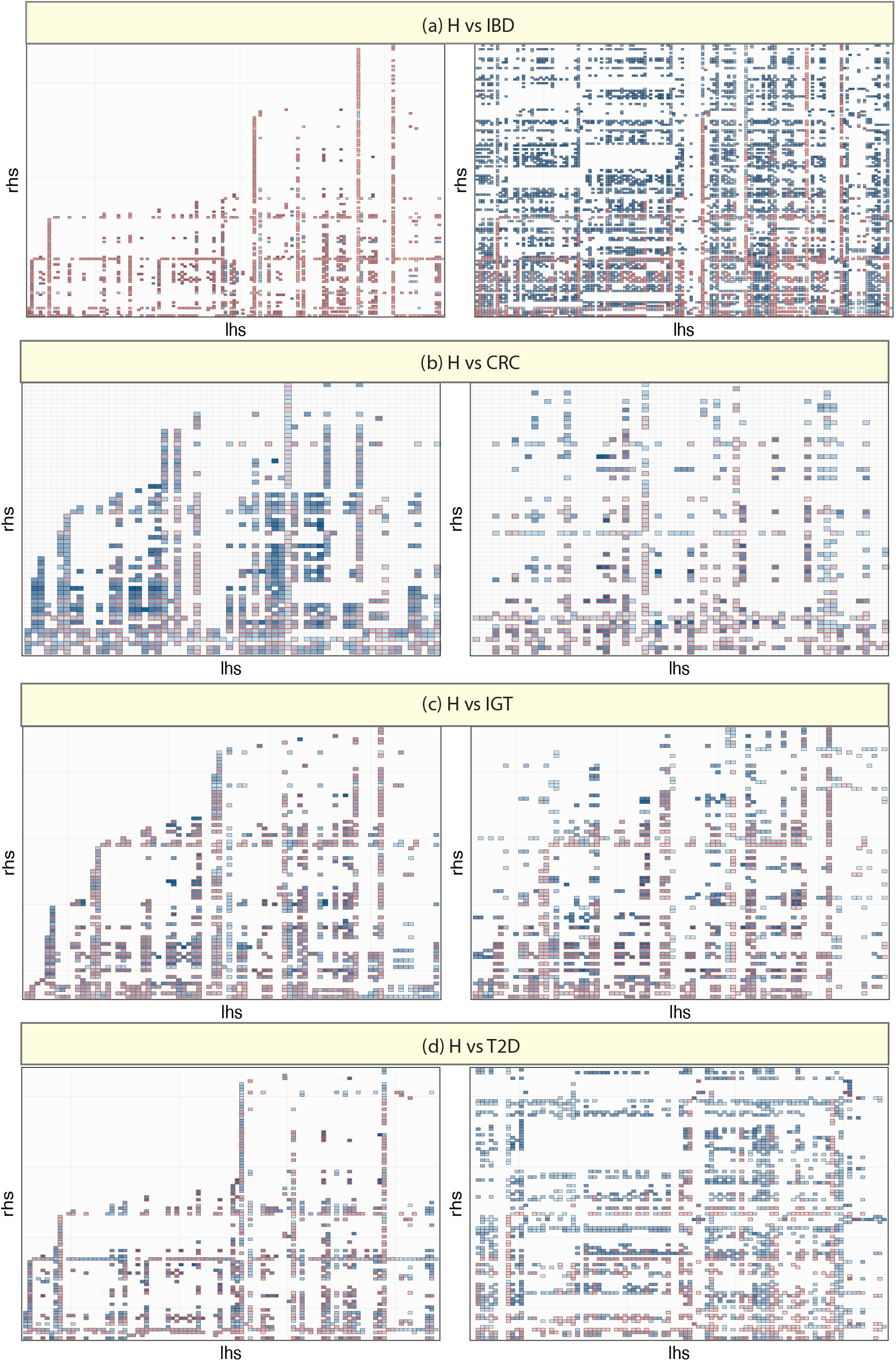
Illustration of the common rules and differentially composed rules inferred from healthy and diseased gut microbiome. Each cell represents an association rule with its LHS itemset on x-axis and its RHS itemset on y-axis. Each cell (rule) is shaded by its *Lift* value. Common rules are highlighted by red boxes.

**Table 1.** Top-20 association rules inferred from healthy gut microbiome samples. This table summarized top-20 association rules (based on their *Lift* values), and their quality measures: support, minimum correlation (minCorr), confidence, lift, and count of occurring in 100 realizations.

| lhs | rhs | support | minCorr | confidence | lift | count |
| --- | --- | --- | --- | --- | --- | --- |
| {F. prausnitzii, D. longicatena, A. hallii} | ⇒ {F. saccharivorans, D. formicigenerans} | 0.705 | 0.174 | 0.942 | 1.197 | 94 |
| {F. saccharivorans, D. formicigenerans} | ⇒ {F. prausnitzii, D. longicatena, A. hallii} | 0.705 | 0.174 | 0.896 | 1.197 | 93 |
| {D. longicatena, A. hallii} | ⇒ {F. prausnitzii, F. saccharivorans, D. formicigenerans} | 0.705 | 0.174 | 0.937 | 1.195 | 91 |
| {F. prausnitzii, F. saccharivorans, D. formicigenerans} | ⇒ {D. longicatena, A. hallii} | 0.705 | 0.174 | 0.898 | 1.195 | 89 |
| {D. longicatena, A. hallii} | ⇒ {F. saccharivorans, D. formicigenerans} | 0.705 | 0.407 | 0.938 | 1.193 | 90 |
| {F. saccharivorans, D. formicigenerans} | ⇒ {D. longicatena, A. hallii} | 0.705 | 0.407 | 0.896 | 1.193 | 90 |
| {F. prausnitzii, F. saccharivorans, A. hallii} | ⇒ {D. longicatena, D. formicigenerans} | 0.705 | 0.174 | 0.893 | 1.191 | 100 |
| {D. longicatena, D. formicigenerans} | ⇒ {F. prausnitzii, F. saccharivorans, A. hallii} | 0.705 | 0.174 | 0.939 | 1.191 | 100 |
| {F. prausnitzii, Anaerostipes hadrus, A. hallii} | ⇒ {F. saccharivorans, D. formicigenerans, B. wexlerae} | 0.703 | 0.260 | 0.885 | 1.191 | 82 |
| {F. saccharivorans, D. formicigenerans, B. wexlerae} | ⇒ {F. prausnitzii, A. hadrus, A. hallii} | 0.703 | 0.260 | 0.945 | 1.189 | 82 |
| {F. prausnitzii, F. saccharivorans, A. hallii} | ⇒ {A. hadrus, D. formicigenerans, B. wexlerae} | 0.703 | 0.260 | 0.891 | 1.189 | 87 |
| {A. hadrus, D. formicigenerans, B. wexlerae} | ⇒ {F. prausnitzii, F. saccharivorans, A. hallii} | 0.703 | 0.260 | 0.938 | 1.188 | 89 |
| {F. saccharivorans, A. hallii} | ⇒ {F. prausnitzii, A. hadrus, D. formicigenerans, B. wexlerae} | 0.703 | 0.260 | 0.886 | 1.188 | 83 |
| {F. prausnitzii, A. hadrus, D. formicigenerans, B. wexlerae} | ⇒ {F. saccharivorans, A. hallii} | 0.703 | 0.260 | 0.942 | 1.188 | 83 |
| {F. saccharivorans, A. hallii} | ⇒ {F. prausnitzii, D. longicatena, D. formicigenerans} | 0.705 | 0.174 | 0.889 | 1.188 | 93 |
| {F. prausnitzii, D. longicatena, D. formicigenerans} | ⇒ {F. saccharivorans, A. hallii} | 0.705 | 0.174 | 0.942 | 1.188 | 93 |
| {A. hadrus, A. hallii} | ⇒ {F. prausnitzii, F. saccharivorans, D. formicigenerans, B. wexlerae} | 0.703 | 0.260 | 0.881 | 1.188 | 85 |
| {F. prausnitzii, D. longicatena, D. formicigenerans, B. wexlerae} | ⇒ {A. hadrus, A. hallii} | 0.703 | 0.260 | 0.947 | 1.188 | 84 |
| {A. hadrus, A. hallii} | ⇒ {F. saccharivorans, D. formicigenerans, B. wexlerae} | 0.704 | 0.270 | 0.883 | 1.187 | 83 |
| {F. saccharivorans, D. formicigenerans, B. wexlerae} | ⇒ {A. hadrus, A. hallii} | 0.704 | 0.270 | 0.946 | 1.187 | 83 |

**Table 2.**
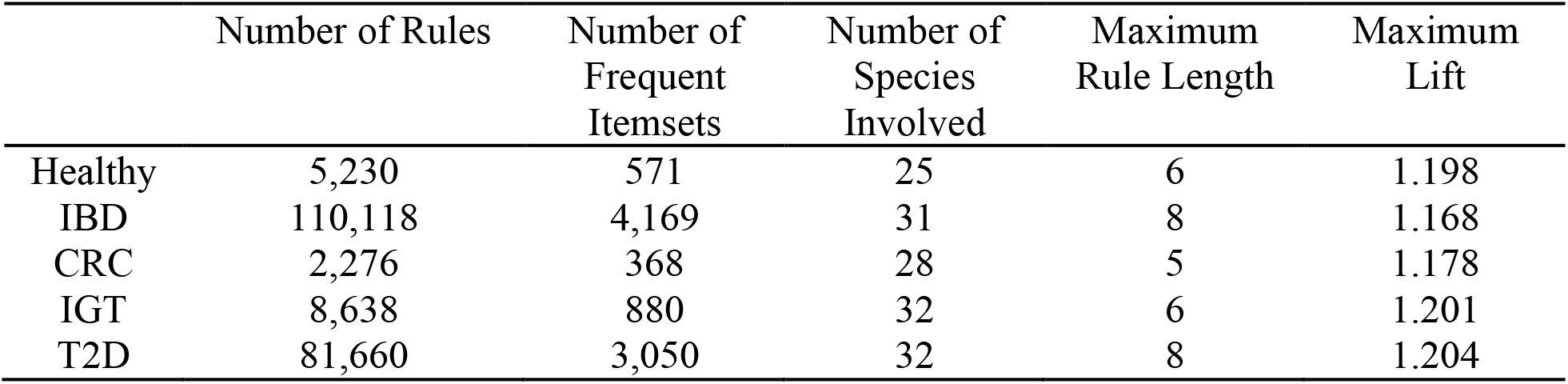
Summary table of association rules inferred from gut microbiome samples with different health/disease status.

**Table 3.** Summary table of comparing association rules inferred from healthy and diseased gut microbiome.

|  | Number of<br>Common<br>Rules | Number of Differentially<br>Composed Rules |  | Number of Different Rules |  |
| --- | --- | --- | --- | --- | --- |
|  |  | From<br>Healthy | From<br>Diseases | Healthy<br>Unique | Disease<br>Unique |
| H (5,230) vs IBD (110,118) | 1,322 | 284 | 1,322 | 3,624<br>(69.3%) | 103,424<br>(93.9%) |
| H (5,230) vs CRC (2,276) | 410 | 660 | 232 | 3,400<br>(65.0 %) | 1,634<br>(71.79%) |
| H (5,230) vs IGT (8,638) | 754 | 456 | 612 | 3,408<br>(65.2%) | 7,272<br>(84.19%) |
| H (5,230) vs T2D (81,660) | 614 | 558 | 2,116 | 4,058<br>(77.6%) | 78,930<br>(96.7%) |

### ARM-enhanced disease classification

Since a large proportion of association rules only occurred in some specific disease status, we can leverage those disease-specific rules to classify diseases. The basic idea is to use ARM as a feature selection tool in microbiome-based disease classification. We employ CPAR (Classification based on Predictive Association Rules) to test this idea. CPAR combines the advantages of both associative classification and traditional rule-based classification to generates a selective set of association rules that differentiate the groups most (Yin & Han, 2003).

As illustrated in **Fig.5a**, given a relative abundance feature table with labeled health/disease status, we randomly sampled half of our samples and apply CPAR with 10-fold on their labeled transaction table to generate a predictive ruleset. A predictive rule is in the form of {*A, B, C*} → *H*, where the LHS contains the presence/absence information of a species set (e.g., *A* and *B* are present, but *C* is absent), and the RHS indicates the health/disease status (e.g., *H* or *D*). We selected rules repetitively occurred in different folds. Those species involved in those predictive rules were selected. Based on their presence or absence status in the predictive rules, the relative abundances of those selected species were assigned signs + or −, and treated as features in downstream classifiers.

**Figure 5.**
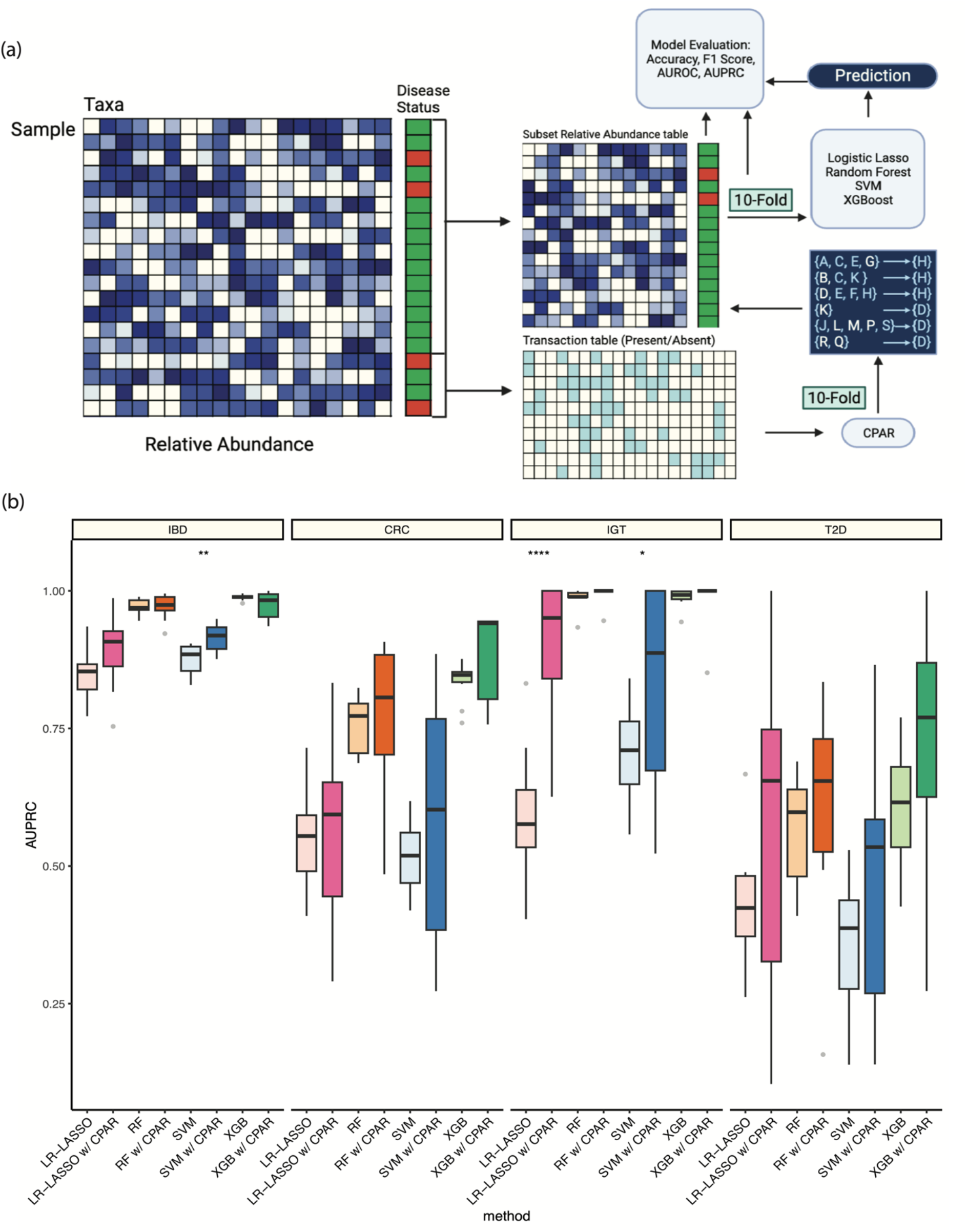
(a) Schematic workflow of classification analysis with ARM-based feature selection. Starting with the relative abundance table with labeled disease status, we randomly sampled half of our samples and apply CPAR with 10-fold on their labeled transaction (present/absent) table to generate a predictive ruleset. The predictive ruleset is similar to the co-occurrence rules before, but it contains the criteria of present (in light-blue) and absent (in white) of a set of taxa in LHS and corresponding disease status in RHS. We selected rules repetitively occurred in different folds, and we selected those features (taxa) included in these rules. Then, the numerical values from our relative abundance table of selected features were extracted and assigned the sign based on the present or absent status in the predictive rules. The selected feature table will then be used to train the models with 10-fold cross validation to access the performance. **(b) Comparative analysis of the classification performance**. The performance of logistic regression model with further feature selection using LASSO, Random Forest (RF), Support Vector Machine (SVM), and eXtreme Gradient Boosting decision trees (XGBoost) on the classification of Inflammatory Bowel Disease (IBD), Colorectal cancer (CRC), Impaired Glucose Tolerance (IGT), and Type 2 Diabetes (T2D). The performance was evaluated by area under the precision-recall curve (AUPRC). The input data of each classifier can be either the whole relative abundance table or the subset relative abundance table by CPAR (denoted as w/CPAR).

In our study, we applied the CPAR-selected features to several standard classifiers: logistic regression model with further feature selection using LASSO (LR-LASSO), Random Forest (RF), Support Vector Machine (SVM), and eXtreme Gradient Boosting decision trees (XGBoost). For each classifier, we compared the classification performance of using the CPAR-selected features, and that of using the original relative abundance table. For the sake of fair comparison, we did not perform any hyperparameter tunning and leave most of them as default settings for each classifier.

For each of the 4 diseases: IBD (n=768), CRC (n=368), IGT (n=199), and T2D (n=164), we used all the 2,815 healthy gut microbiome samples in CMD as the healthy controls (n=2,815). Given the highly unbalanced data for each disease classification task, we chose the area under the precision-recall curve (AUPRC) as the performance metric. We performed 10-fold cross-validation to evaluate the classification performance in terms of AUPRC. Our results are shown in **Fig.5b**. After the feature selection with CPAR, we usually get 100-130 species selected, rendering a significant feature reduction comparing to 200 species from the original relative abundance table. In terms of the classification performance, we observed a noticeable improvement for all the 4 diseases and all the 4 classifiers.

## DISCUSSION

We performed ARM for human gut microbiome samples in a large-scale curated metagenomic database. Several modifications on the traditional ARM were implemented to accommodate the nature of microbiome data. We observed both common and disease-specific association rules. The latter further helped us to perform feature selection and enhance microbiome-based disease classification.

In the top association rule (with the highest *Lift*) minedfrom healthy gut microbiome samples: {*F. prausnitzii, D. longicatena, A. hallii*} }⟹ {*F. saccharivorans, D. formicigenerans*}, all the five species are in the phylum of Firmicutes. The co-occurrence of those species is also widely observed among the frequent itemsets generated from diseased samples as shown in Figure 3. It has been demonstrated that the main butyrate producing-bacteria in the human gut belong to the phylum Firmicutes (Parada Venegas et al., 2019). Indeed, *Faecalibacterium prausnitzii, Anaerobutyricum hallii, Fusicatenibacter saccharivorans* were found to be butyrate producers(Deleu et al., 2021; Xie et al., 2021). *Dorea* genus was also reported to be associated with the butyrate level (Fu et al., 2019). Butyrate has beneficial effects on both cellular energy metabolism and intestinal homeostasis as a least produced short chain fatty acids, but it is the major energy source for colonocytes (H. Liu et al., 2018).

ARM-based feature selection holds great promise in microbiome-based disease classification. As demonstrated in the performance of four widely used classifiers on four diseases, the improvement of the classification performance is significant (especially in the case of IGT and T2D, which suffer from the data imbalance issue the most). Further improvements of the microbiome-based classification can be achieved by systematically tunning the hyper-parameters.

Extracting biologically meaningful information from association rules is still a demanding task. From the association rules inferred from diseased microbiome samples, systematic approaches are needed to interpret the disease-specific rules and understand the underlying mechanism.

